# Concern noted: A descriptive study of editorial expressions of concern in PubMed and PubMed Central

**DOI:** 10.1101/106757

**Authors:** Melissa Vaught, Diana C. Jordan, Hilda Bastian

## Abstract

**Background:** An editorial expression of concern (EEoC) is issued by editors or publishers to draw attention to potential problems in a publication, without itself constituting a retraction or correction.

**Methods:** We searched PubMed, PubMed Central (PMC), and Google Scholar to identify EEoCs issued for publications in PubMed and PMC up to 22 August 2016. We also searched the archives of the Retraction Watch blog, some journal and publisher websites, and studies of EEoCs. In addition, we searched for retractions of EEoCs and affected articles in PubMed up to 8 December 2016. We analyzed overall historical trends, as well as reported reasons and subsequent editorial actions related to EEoCs issued between August 2014 and August 2016.

**Results:** After screening 5,076 records, we identified 230 EEoCs that affect 300 publications indexed in PubMed, the earliest issued in 1985. Half of the primary EEoCs were issued between 2014 and 2016 (52%). We found evidence of some EEoCs that had been removed by the publisher without leaving a record and some were not submitted for PubMed or PMC indexing. A minority of publications affected by EEoCs had been retracted by early December 2016 (25%). For the subset of 92 EEoCs issued between August 2014 and August 2016, affecting 99 publications, the rate of retraction was similar (29%). The majority of EEoCs were issued because of concerns with validity of data, methods, or interpretation of the publication (68%), and 31% of cases remained open. Issues with images were raised in 40% of affected publications. Ongoing monitoring after the study identified another 17 EEoCs to year’s end in 2016, increasing the number of EEoCs to 247 and publications in PubMed known to be affected by EEoCs to 320 at the end of 2016.

**Conclusions:** EEoCs have been rare publishing events in the biomedical literature, but their use has been increasing. Most have not led to retractions, and many remain unresolved. Lack of prominence and inconsistencies in management of EEoCs reduce the ability of these notices to alert the scientific community to potentially serious problems in publications. EEoCs will be made identifiable in PubMed in 2017.

## Background

An editorial expression of concern (EEoC) enables journal editors or publishers to quickly indicate to readers that they have a “concern about the integrity of a published article” [1]. The Council of Science Editors (CSE) says an EEoC aims to draw attention to problems in a publication, “but it does not go so far as to retract or correct an article” [2]. The term was formally introduced as a standard for biomedical journals in 1997 by the International Committee of Medical Journal Editors (ICMJE) [3], although some other terms are used for the same purpose, such as “notice of concern” or “publisher’s note”.

As with retractions, Council on Publication Ethics (COPE) advises that author consent is not required for issuing an EEoC. COPE’s guidelines encourage editors to consider an EEoC if [4]:

- “they receive inconclusive evidence of research or publication misconduct by the authors
- there is evidence that the findings are unreliable but the authors’ institution will not investigate the case
- they believe that an investigation into alleged misconduct related to the publication either has not been, or would not be, fair and impartial or conclusive
- an investigation is underway but a judgement will not be available for a considerable time”.

There is relative consensus between the advice from publisher and editor organizations on dealing with issues of research integrity. However, considerable differences in policy and practice remain between journals. The policies of several journals and publishers in relation to EEoCs refer to, or quote, the COPE guidelines, although one publisher qualifies the level of concern required to issue an EEoC as “well-founded suspicions of misconduct” [5].

Journal editor Ana Marusic and her colleagues wrote that some editors are concerned about their capacity and/or legal position in investigating allegations of scientific misconduct. They may refer concerns to authorities, but there may be no, or only an unsatisfactory, response. Marusic et al also pointed to the lack of adverse consequences for editors when they avoid dealing with research integrity cases [6].

Another journal editor, Richard Smith, contended that journals “cannot provide due process”, to which anyone faced with a serious accusation has a right [7]. Chris Graf et al provided one major publisher’s perspective on the role of editors in publication ethics. Editorial adjudication was supported, and the importance of opportunities for author comment and appeal noted [8].

There is some legal precedent on the use of EEoCs in the United States. In 2015, an author filed an application with a U.S. court that, if successful, would have required the American Diabetes Association to remove four EEoCs from their journal, *Diabetes.* The Court dismissed that application, however the author pursued defamation action as well [9]. That motion was also dismissed, with the Court finding that, “the Expression of Concern is a statement of opinion that is not actionable for defamation” [10].

Although there have been many studies of retractions, which are tagged in key biomedical literature databases, we identified only three studies of the extent of use of EEoCs. Noonan and Parrish discussed the positions of various medical journal editors in 2008, and identified 15 EEoCs in medical journals, plus one in an engineering journal [11]. Grieneisen and Zhang searched for EEoCs across the broad science and biomedicine literature in mid-2011. They identified EEoCs affecting 58 publications between 2000 and 2011 across 30 journals, with 70% occurring since 2008 [12]. An analysis by Roig was available as a conference abstract only [13]. Roig identified 95 EEoCs in the PubMed database up to May 2015, affecting 124 articles.

PubMed is a free public database comprising more than 26 million citations for biomedical literature, operated by the U.S. National Library of Medicine (NLM) at the National Institutes of Health (NIH). The majority of the PubMed database consists of the MEDLINE journal database. PMC, formerly PubMed Central, is the NLM’s free full-text archive of biomedical literature [14].

Although EEoCs are not as definitive as retractions, the seriousness of their content justifies ensuring that they are prominent in PubMed’s databases. However, they are not tagged with a MEDLINE publication type, as retractions have been since 1984 [1]. PMC began tagging EEoCs as an article type, within PMC only in 2013 [15].

We undertook a systematic search to identify EEoCs affecting publications in PubMed and PMC to understand the range of publishing practice, and assess the value and feasibility of tagging these notices.

## Methods

### Eligibility criteria

EEoCs and publications in any language were eligible for inclusion.

1. Editorial expression of concern (EEoC) A notification of concern about a publication was included as an EEoC if it was attached to, or directly referencing, a publication that was indexed in PubMed or PMC and was:

- Authored by one or more editors, or by the publishers of a journal; and
- Published in the same journal as the article, or in another of the publisher’s active journals where a journal has ceased publication. The EEoC did not have to be independently indexed in PubMed to be included. EEoCs which were no longer available at PubMed, PMC, or the journal website, but could be confirmed at a reliable source, were also included. An additional EEoC or follow-up note to an EEoC reporting on developments in an investigation that did not explicitly retract the concern was included as an additional EEoC. Letters to the editor called “expression of concern” were excluded, as were EEoCs identified for publications that were not indexed in PubMed.
2. Retracted EEoC Formal notices by editors or publishers withdrawing or retracting an EEoC were included as EEoC retractions.
3. Affected publication Publications which were the subject of one or more EEoCs were included if they were the direct subject of editorial (or publisher) concern, and they were indexed in PubMed. Publications referred to in an EEoC that were not themselves the subject of concern were excluded, for example, a publication that the affected publication may have duplicated or plagiarized.

### Search strategy for EEoCs

Details of search strategies are reported in accordance with PRISMA reporting guidelines for systematic reviews [16] in Additional File 1, with data on searches and screening in Additional File 2. The search strategies were developed by all three authors. The strategies were not peer reviewed.

Using search terms including “expression of concern”, “notice of concern”, “note of concern”, and “statement of concern”, we searched PubMed and PMC to identify EEoCs issued for publications indexed in PubMed. We distinguished withdrawn EEoCs from retractions of publications that were the subject of EEoCs. All PubMed and PMC searches were done by a single author (DJ or MV), and screened by one author (HB) and a second person (MV or another colleague). Rayyan was used for the bulk of screening PubMed search results [17]. Searches for EEoCs were finally updated on 22 August 2016, with the final update of search for retractions of EEoCs on 7 December.

Between August and December, we undertook additional searches for EEoCs issued up to 22 August 2016. We searched Google Scholar, the tagged “expression of concern” archive of the Retraction Watch blog [18], and 9 publisher websites for EEoCs. We hand-searched for references to EEoCs in studies of EEoCs and the CSE’s white paper on publication ethics [2].

For all journals where an EEoC was identified that had not been submitted to PubMed or PMC or had not been found by searching PubMed/PMC, the journal website was searched with the terms used in the EEoC(s) which had been identified. This resulted in searches of 17 journals.

Searches and pre-screening of Google Scholar, publisher, and journal websites were done online by a single author (HB or MV). Data from Retraction Watch posts was extracted by one author (DJ), and posts were screened by two authors (DJ, MV).

All differences in selection of records included in dual-screening were resolved by discussion. The final inclusion of all EEoCs, retracted EEoCs, and affected publications were agreed by two authors (HB, MV). Included and excluded records are available in Additional File 3. On 6 December, each EEoC in the final database which had not been indexed in PubMed was searched for in PubMed.

### Data collection and analysis

For all EEoCs and affected publications, the following data were extracted from the MEDLINE record or the journal website where an EEoC was not PubMed-indexed:

- Date of EEoC;
- Date of retraction for any retracted EEoC;
- Date of publication for affected publications;
- Date of retraction for any retracted or corrected and republished affected publication;
- Journal.

Data were extracted from the MEDLINE records by one author (DJ), with data from journal websites checked by two authors (DJ, MV). Retraction or correction and republication of affected publications in PubMed was identified by searching for those tagged publication types “retracted publication” and “corrected and republished article”, confirmed by two authors (DJ, MV). In addition, all dual-confirmed retractions and corrected republished articles identified by the authors in a companion ongoing project were also searched, and additions confirmed by two authors (HB, DJ). A final update search for subsequent retractions and corrected republished articles was done on 8 December 2016. Linkages between EEoCs, affected publications, and post-EEoC events are available in Additional File 4.

Analyses were performed using RStudio 1.04 running R 3.3.1 [19, 20]. Data import and processing were supported by the packages openxlsx, plyr, dplyr, tidyr, and stringr [21, 22, 23, 24, 25]. Graphs were created using ggplot2, survival, survminer, and riverplot [26, 27, 28, 29, 30]. Data for this project are deposited at the Open Science Framework [31].

Summary statistics were used to describe the cohort. Data on number of affected publications by journal, and number of EEoCs and affected publications by year are available in Additional File 5. We assessed survival of articles to EEoC via Kaplan-Meier analysis, using the survfit function in the survival package for R. Summary statistics were calculated for time from EEoC to retraction, where applicable. Dates for PubMed record creation (CRDT) were used to calculate times between publication and EEoC, and between EEoC and retraction of affected publications. We assumed that, for a given journal, the time lag between publishing and indexing is similar for all publication types. Data for these analyses are available in Additional File 6.

We also undertook an analysis of reported reasons for concern, and subsequent corrections and retractions of publications, for EEoCs issued between August 2014 and August 2016. For all affected publications, the journal website was examined by two authors to identify any additional editorial actions that were not submitted to PubMed.

The categories for reasons for EEoCs were iteratively developed during coding by HB and MV, starting from criteria used in Decullier [32] and the definitions for research misconduct by the HHS Office of Research Integrity [33].

The following fields were classified by two authors (HB, MV), with disagreements resolved by discussion:

- Reason for expressing editorial concern and subsequent action;
- Whether or not images, for example immunoblot or microscopy images, were the, or a, reason for the EEoC or subsequent action;
- Current status of the EEoC;
- Who had adjudicated in cases of findings of misconduct;
- Whether or not the case was investigated by authors’ employer or funding agency, and if so, whether or not the case had been referred by or to the journal.

A detailed description of definitions and a key for coding of the content analysis are available in Additional File 7, with the data in Additional File 8.

## Results

We retrieved or pre-screened online 5,076 records, finally dual-screening 1,208 unique records. We identified 230 EEoCs that affect 300 publications indexed in PubMed (Figure 1 and Additional File 3).

**Figure 1.**
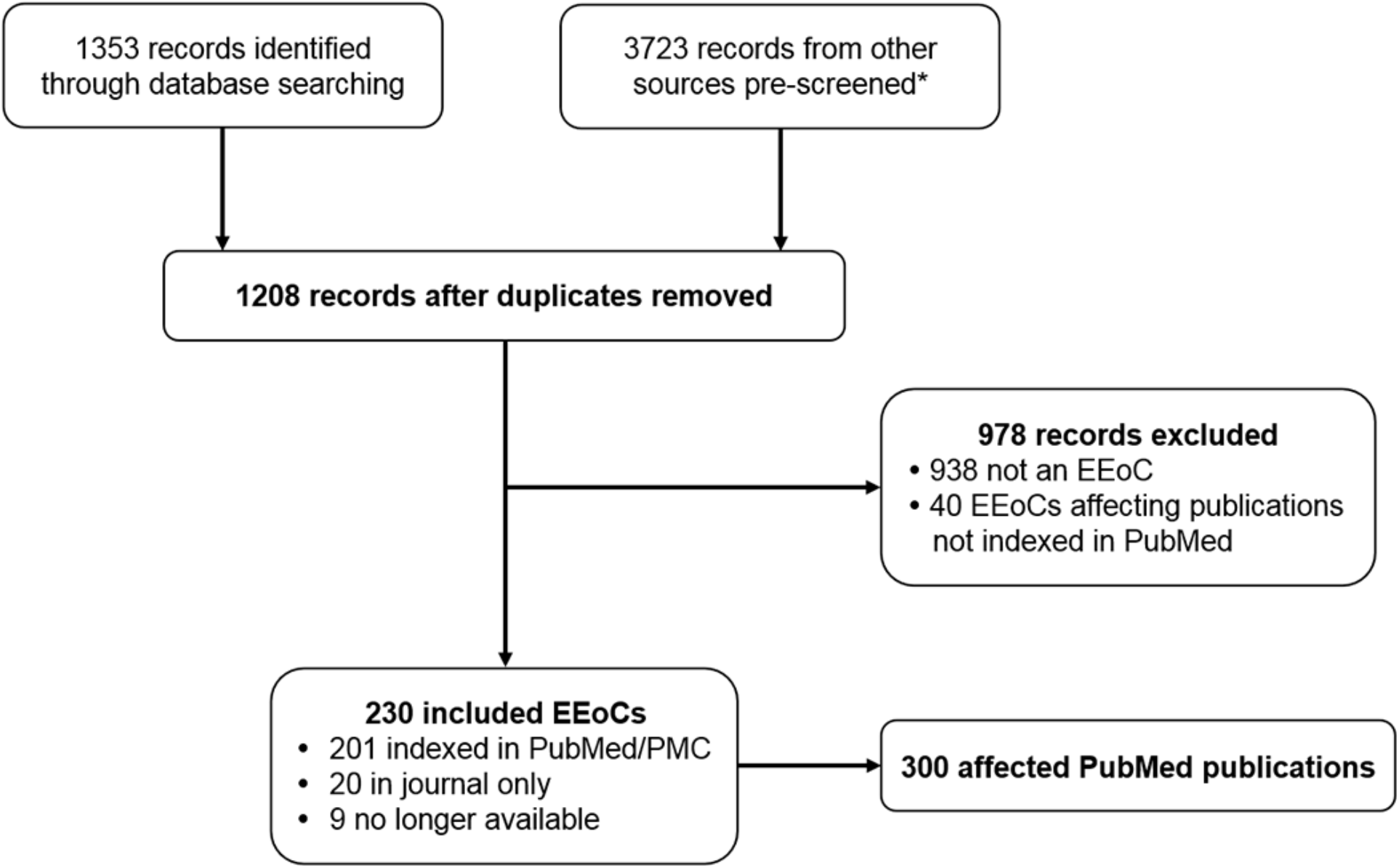
Search results for expressions of concern (EEoCs) and affected publications. *Records could not be downloaded from Google Scholar, and those results along with other online sources, were prescreened online. Only likely EEoCs were added to the records for formal dual screening.

We excluded 40 EEoCs affecting publications not indexed in PubMed, including one that affected the complete issue of a journal. Those excluded EEoCs, along with the identifiers for excluded records from the PubMed and PMC searches, are included in Additional File 3. We encountered only 3 non-English records in our searches, all of which were clearly not EEoCs.

The majority (87%) of EEoCs were individually indexed in PubMed/PMC. Of the remaining 29, 9 are no longer available online at the journal. The EEoCs which had not been indexed at PubMed or PMC affected 13 journals, and 2 of those journals had both submitted and unsubmitted EEoCs.

Of the publications affected by the 9 EEoCs which were no longer available, 5 were subsequently retracted, 2 were followed by publisher/editor statements, 1 was formally withdrawn, and 1 article was replaced by a version with a correction.

Evidence of the 9 EEoCs which were no longer available were found in a retraction notice (1), the retraction of an EEoC (1), follow-up statements by publishers (2), reported in a news piece in a journal (1), or copied in full at Retraction Watch (4). In addition, 1 EEoC submitted to PubMed was subsequently withdrawn without a formal retraction notice: the EEoC at the publisher site had been replaced with the withdrawal notice.

In total, there were 221 primary EEoCs, of which 6 had been formally retracted (a notice explicitly referring to withdrawal or retraction), with 9 follow-up EEoCs (Table 1).

**Table 1.**
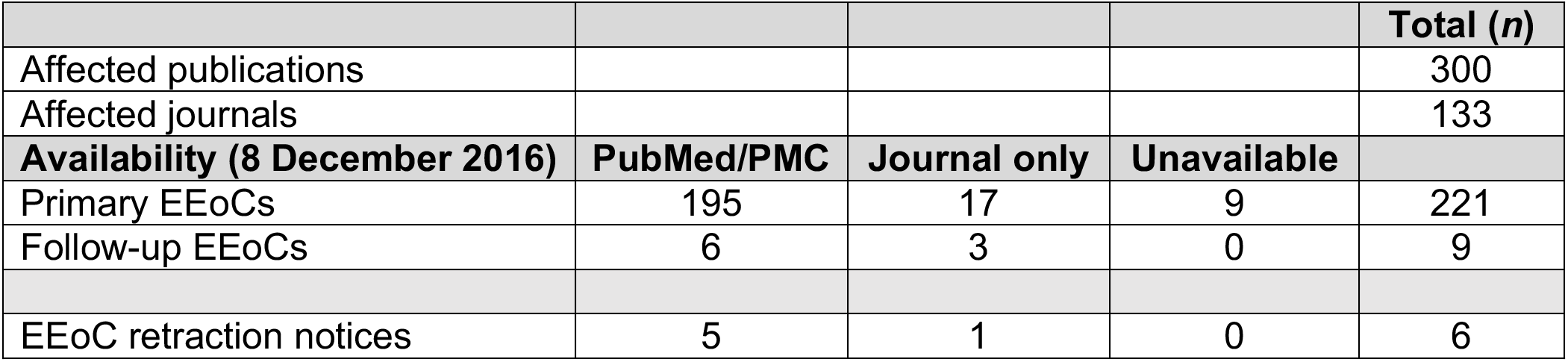
PubMed/PMC-indexed publications and journals affected by EEoCs, and EEoCs by type and availability.

Publications affected by known EEoCs were found in 133 journals (Additional File 5). Most of those journals had a single affected publication (n = 91, 68%), with 2 to 4 affected publications in 27 journals, and 5 or more affected publications in 15 journals. The highest number of affected publications for a single journal was 41, where the publisher issued an EEoC in 2014 for all publications in a six-month period in 2012.

The first EEoC found was from 1985, although the first explicit use of the phrase “expression of concern” was not until 2000. EEoCs began to appear more frequently in 2005, increasing further in the past 5 years (Figure 2). The highest number in a year was in 2016, an incomplete year in this dataset. Half the primary EEoCs were issued between 2014 and 2016 (52%).

**Figure 2.**
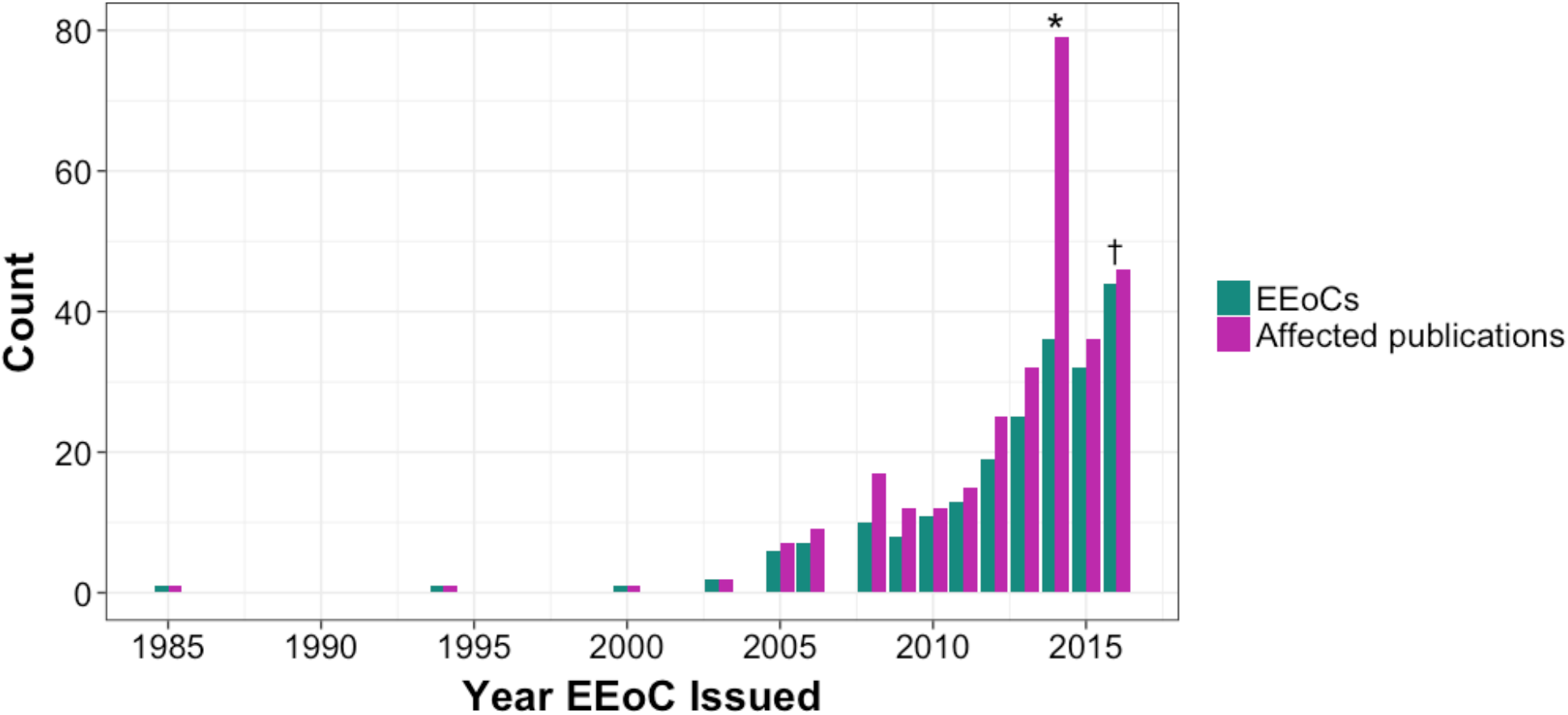
Number of primary EEoCs and affected publications by year EEoC issued (1985-2016). * A single EEoC was issued for 41 publications in a 6-month period for a journal by its publisher. † Incomplete year (data collected to 22 August 2016). Subsequent monitoring increased the number of EEoCs in 2016 to 59, affecting 66 publications (post-study data not included in figure). Only primary EEoCs are included. Year of issue could not be definitely assigned for 5 EEoCs. Based on information from the associated Retraction Watch post, 3 of these were published in late 2014 or early 2015. The other 2 were added as online notes, with no date of posting.

After the study was completed, we identified a further 17 EEoCs to year’s end in 2016 by monitoring new PubMed and PMC entries with simplified searches, as well as Retraction Watch. This brought the total number of EEoCs to 247 and increased the number of publications in PubMed known to be affected by EEoCs to 320 at the end of 2016. One of those studies, identified via Retraction Watch, occurred within the study period. However, none of those additional 17 EEoCs or affected publications are included in any of the study’s analyses. Data for those EEoCs and affected publications are available in Additional File 9.

Publications usually continue to stand following an EEoC, and EEoCs themselves are rarely retracted or receive follow-up notices (Figure 3). However, 25% of the publications affected by EEoCs had been retracted as of 8 December 2016, 1 of which was corrected and republished. Two additional affected publications were corrected and republished, but without being formally retracted.

**Figure 3.**
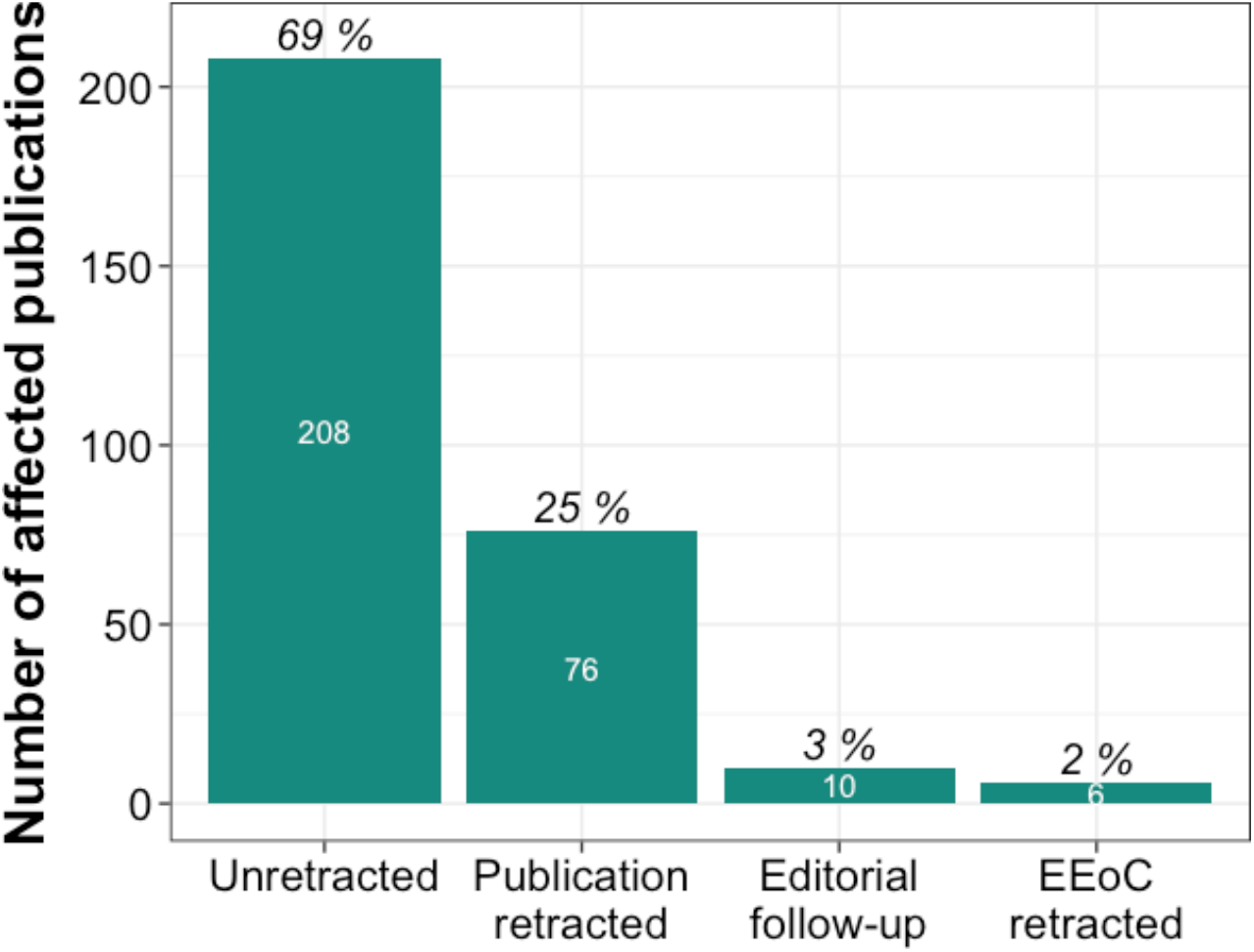
Affected publications, retracted publications, and follow-up and retracted EEoCs.

We ran survival analyses for the 260 affected publications (87%) which had unique PubMed records available for the publication as well as EEoC. Although some EEoCs were issued within days of publication, months or years had typically passed (Figure 4). EEoCs were issued for 83 affected publications (32%) after more than 5 years, and after more than 10 years for 25 (10%).

**Figure 4.**
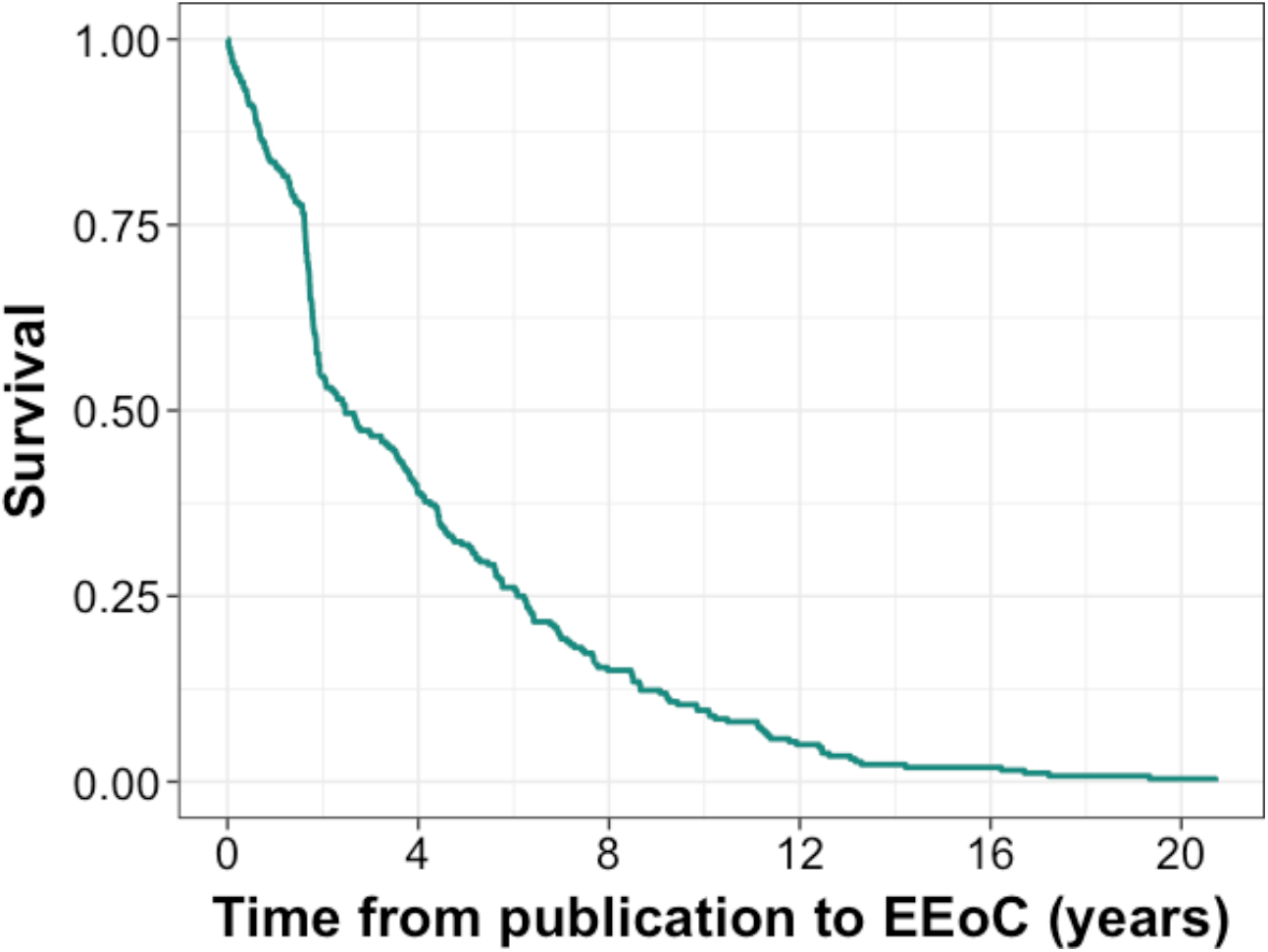
Time from publication to EEoC in years (n = 260).

The mean time from publication to original EEoC was 1,516±1,415 days, and the median was 900 days with an IQR of 1,640. The longest time between publication and an EEoC was more than 20 years.

Of the 260 affected publications included in survival analyses, 64 (25%) had been retracted as of 8 December 2016. Survival to retraction could not be calculated for 2 of these publications because unique records were not available for the retraction. The 62 publications are 82% of all retracted publications shown in Figure 3.

More than half of these retractions occurred within the year after the primary EEoC was issued (*n* = 38, 61%), and 57 publications (92%) were retracted within 2 years of primary EEoC. The mean time from EEoC to retraction was 299±245 days, and the median was 263 days with an IQR of 333. The longest gap between original EEoC and retraction was just under 3 years.

## Analysis of 2014-2016 sample

There were 92 EEoCs issued between August 2014 and August 2016, affecting 99 publications. The rates of retraction (29%), follow-up EEoC (4%), and retraction of EEoC (1%) were similar to those for the full sample. In addition, 6% of affected publications had subsequent errata.

Seven EEoCs were no longer available at the publisher site, PubMed or PMC, because they were over-written or removed without formal retraction. Where EEoC text was available at Retraction Watch, it was included. Only 2 EEoCs could not be coded, affecting 2 publications. We evaluated the other EEoCs, along with all subsequent follow-up EEoCs, corrections (errata or corrected and republished articles), and retractions for those records from 8 December for the final analysis.

Concerns about validity of data, methods, or interpretation were expressed for 66 publications (68%) (Table 2). Allegations or findings by others of research misconduct were noted for 11 of publications (11%). Of those, where stated, the allegation or finding was on the part of the journal for 4 publications and an external adjudication for 3. Research misconduct included fabrication and/or falsification, plagiarism, scientific misconduct that was not specified, or ethical misconduct. (For more details of the classifications see Additional File 7, and for content analysis data see Additional File 8.)

**Table 2.**
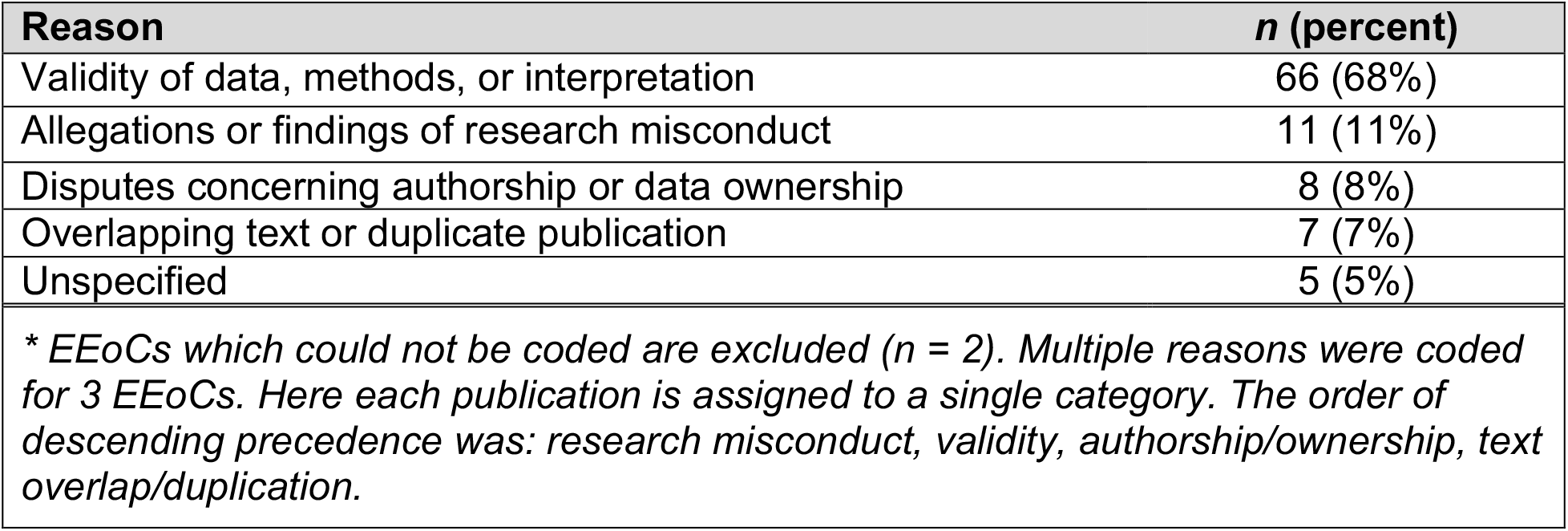
Reasons for EEoC about publications (n = 97*) (August 2014 – August 2016).

EEoCs appeared to represent the endpoint of editorial investigation for 28 publications (28%). This occurred, for example, when concerns had been addressed without affecting the publication, or when concerns were raised about data validity, but primary data were no longer available for review. In one instance where an EEoC served as final notice to readers, an erratum was published at the same time.

An additional 37 cases were closed by subsequent action: 29 publications were retracted, 5 were corrected, and 3 cases were finalized with follow-up EEoCs. We found current status ambiguous for 2 publications, and considered the status of the 30 remaining cases open (31%).

Reasons for actions were sometimes more specific or differed in other ways in subsequent notices where the text for the action was available (Table 3). Allegations or findings by others of research misconduct were noted for 15 of publications (38%). Of those, where stated, the allegation or finding was on the part of the journal for 3 publications, an external adjudication for 10, and both journal and external adjudication for 1.

**Table 3.** Reasons noted in subsequent actions (n = 40*) (August 2014 – August 2016).

| Reason | n (percent) |
| --- | --- |
| Validity of data, methods, or interpretation | 20 (50%) |
| Allegations or findings of research misconduct | 15 (38%) |
| Disputes concerning authorship or data ownership | 1 (3%) |
| Other | 1 (3%) |
| Unspecified | 3 (8%) |
| *\* Includes 2 publications where the EEOC was unavailable for coding, but subsequent action was coded. Multiple codes were assigned for 7 actions. Here each publication is assigned to a single category. The order of descending precedence was: research misconduct, validity, authorship/ownership. If multiple subsequent events occurred, the coding for the final event is reported.* |  |

To show how classifications changed, we paired and graphically displayed classifications for the primary EEoC and the subsequent notice (Figure 5). In particular, unspecified concerns about validity in EEoCs were later specified as research misconduct in 9 publications.

**Figure 5.**
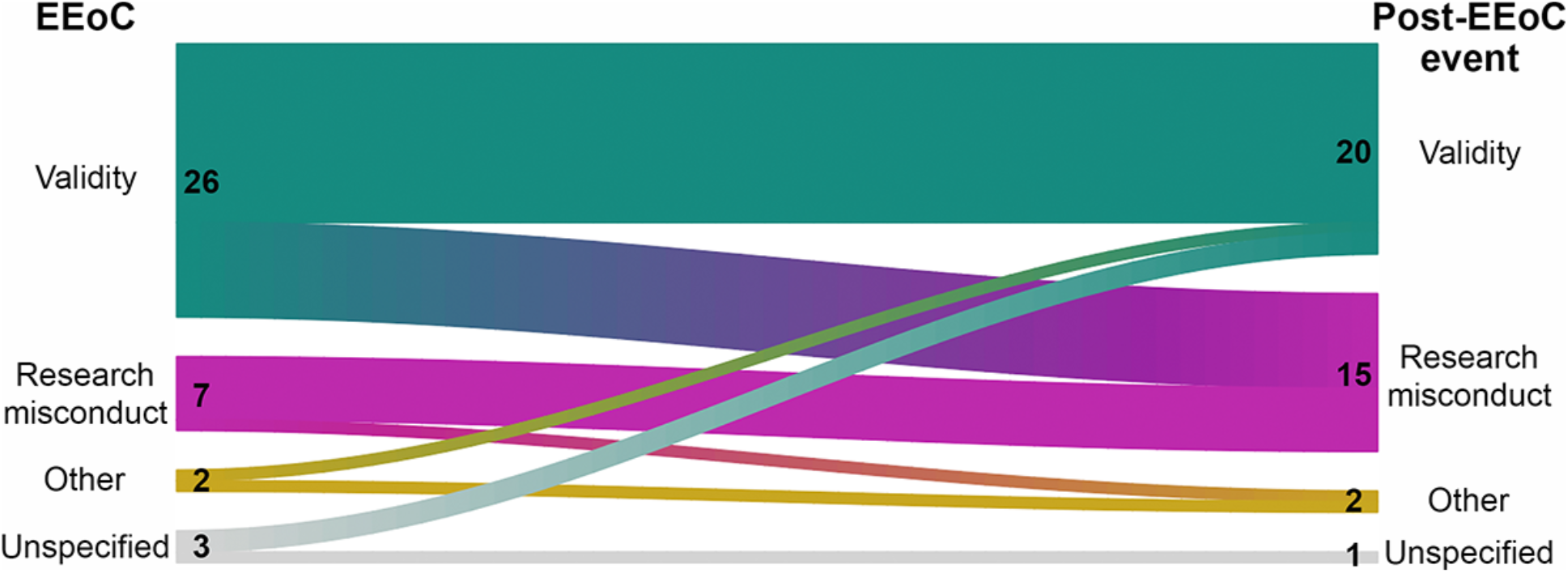
Reasons for EEoC about publication versus reasons noted in subsequent actions.

Involvement of the authors’ employer or funding agency either during the investigation, or because of referral by the journal to them, was reported in EEoCs for 46 out of 97 publications. For 14 publications (14%), the matter had apparently been referred to the journal by the institution. Journals referred issues to authors’ institutions for further investigation in 30 cases (31%). In 2 cases, both journals and institutions were investigating, with the order unclear. For the remaining 51 publications (53%), referral of issues was not stated or not applicable in the subsequent action. In subsequent actions, involvement of the authors’ employer or funding agency was reported for a further 4 publications.

Images, for example, immunoblots or microscopy images, were noted as a cause for the primary EEoC in 29 publications (30%). Issues with images were noted for an additional 9 publications in retraction notices, and 1 publication in an erratum. Thus, image issues were raised in 40% of publications affected by EEoCs.

## Discussion

EEoCs are a rare but growing class of publishing events in the biomedical literature. They usually relate to concerns about validity of data, methods, or interpretation in publications, often years after publication. An EEoC is often the endpoint for a publication, and when there is further action, retraction is more common than correction.

Although a substantial proportion of EEoCs resulted in retractions, most had not at the time our study concluded in early December 2016. However, a substantial proportion of EEoCs have been issued in the last few years. As we estimate that 31% of publications affected by EEoCs in the last 2 years remain unresolved, the retraction rate for much of this cohort of publications may increase.

Our search for EEoCs was extensive, but we remain unsure about how many EEoCs have been issued. We identified some EEoCs that journals had not submitted to PubMed or PMC, but only searched some individual journal and publisher websites. All EEoCs are not readily identifiable via Google Scholar, and EEoCs are sometimes removed from publisher websites or replaced with other notices. Although we had no language restrictions, our search strategies would not have identified an EEoC entirely in a language other than English. Although some EEoCs were given the same titles as the affected publication with no additional specification, we did not screen all records in PubMed with identical titles in the same journals.

EEoCs are an important mechanism for timely alerts to the scientific community of potentially serious problems in the literature. Drawing attention more quickly and effectively to affected publications is an essential first step to reducing the lengthy persistence of known error in the biomedical literature [34, 35, 36, 37]. However, EEoCs have been issued by only a small proportion of the thousands [38] of currently active biomedical journals.

Editor and publisher assignment of EEoCs is broadly consistent with guidelines, although labelling and content management vary greatly. We experienced a variety of difficulties in identifying EEoCs. In many journals, categories of unusual publishing events like retractions and EEoCs appear to be unplanned for within the publishing system.

EEoCs can have a variety of titles other than “expression of concern”, and do not necessarily include the title of the affected publication as recommended by ICMJE [39]. They can appear within journal sections dedicated to corrections, retractions, or letters, in supplementary information, or as text inserted with no specified record created. Consequently, these notices can escape both indexing and library services, despite their importance for users of the literature. Inserting EEoCs into fields titling them as retractions or another category also sometimes makes classifying these notices complex. Practice in relation to retraction of EEoC also varies.

Some of the titles used may make it difficult for literature users to appreciate the significance of the notice. The notices are often not prominently placed and thus unlikely to be seen. Some journals facilitate access by including the full text of the notice within the abstract field for the record, but EEoCs can also be behind paywalls.

The ICMJE recommends ensuring proper indexing for both EEoCs and retractions [39] and the CSE recommends the assignment of a DOI [2]. Because this practice is not routinely followed and EEoCs do not always remain available, the historical record of publications is not complete. There is some ambiguity in COPE’s guidelines on this. COPE states that if a retraction confirms the concern about the publication, “the expression of concern should be replaced by a notice of retraction”, implying removal of the original concern from the scientific record is accepted practice. If investigation clears the concern, on the other hand, the publication should receive “an exonerating statement linked to the expression of concern” [4].

Clarity and consistency in recommendations for managing EEoCs is needed. It would be helpful for users if all EEoCs and follow-up notices to EEoCs were clearly identified as either expressions or notices of editorial concern to distinguish them from other communications to readers. Where editorial concern is resolved, EEoCs can be retracted in the same way as a publication. To ensure the integrity of the historical record about publications, as well as to improve the ability of EEoCs to alert readers, permanent free-standing records need to be created for EEoCs, linked prominently to the affected publication.

We established that ongoing screening of new records in PubMed and PMC with limited search fields, as well as Retraction Watch, is not onerous, and is likely to capture most if not all EEoCs. Together with the new capability for publishers to tag and link affected publications within the PubMed Data Management system, this will enable more timely linkage of EEoCs and affected publications in PubMed [40, 41]. EEoCs will be made prominently identifiable to PubMed users in 2017. The classification of EEoCs here may not, however, represent final decisions for PubMed/PMC.

## Conclusions

EEoCs have been rare publishing events, but their use is increasing in the biomedical literature. Editorial use of EEoCs is broadly in line with the purpose described in relevant guidelines, but their labelling, management, display, and submission for library indexing is inconsistent, as are the recommendations of relevant guidelines. This reduces the ability of these notices to alert the scientific community to potentially serious problems in publications. Most EEoCs have not led to retractions or corrections, and many remain unresolved.

## Declarations

### Acknowledgements

–The views expressed by the authors are personal, and do not necessarily state or reflect those of the National Institutes of Health or the U.S. Government. We gratefully acknowledge the assistance of Valerie Virta, who participated in some of the screening of PubMed/PMC search results.

### Authors’ contributions

–All authors participated in the design of this study, screening of PubMed and PMC records, and preparation of the manuscript. MV and HB classified the content of expressions of concern. MV and DJ managed the data and prepared the figures. MV and HB are the guarantors for the data and contents.

### Consent for publication

–Not applicable.

### Ethics approval and consent to participate

–Not applicable.

## List of abbreviations

COPE: Council on Publication Ethics
CSE: Council of Science Editors
DOI: Digital object identifier
EEoC: Editorial expression of concern
ICMJE: International Committee of Medical Journal Editors
NCBI: National Center for Biotechnology Information
NLM: U.S. National Library of Medicine
PMC: PubMed Central
PMID: PubMed identification number
PRISMA: Preferred Reporting Items for Systematic Reviews and Meta-Analyses

## Additional files

Additional files available at: https://osf.io/8xbqy/

Additional File 1: Search strategies
Additional File 2: Results of individual search strategies
Additional File 3: Included and excluded records
Additional File 4: Linkages between EEoCs, affected publications, and post-EEoC events
Additional File 5: Number of affected publications by journal and number of EEoCs and affected publications by year
Additional File 6: Publication survival time to EEoC and retraction
Additional File 7: Definitions and key for content analysis of EEoCs and post-EEoC events
Additional File 8: Content analysis of EEoCs and post-EEoC events
Additional File 9: EEoCs in 2016 identified in routine monitoring post-study

